# Modeling Western Pacific Amyotrophic Lateral Sclerosis and Parkinsonism–dementia Complex with Microglia Containing Cerebral Organoids Derived from Induced Pluripotent Stem Cells

**DOI:** 10.1101/2021.08.06.455467

**Authors:** Yiling Hong, Xu Dong, Lawrence Chang, Mariann Chang, Chen Xie, Jose S. Aguilar, Qingshun Q. Li

## Abstract

Western Pacific Amyotrophic Lateral Sclerosis and Parkinsonism–dementia Complex (ALS-PDC) is a neurodegenerative disease linked to the traditional consumption of cycad seeds by the Chamorro people of Guam. Little is known about the etiological role of cycad toxin in ALS-PDC. Patient-derived induced pluripotent stem cells were derived from age- and sex-matched affected and unaffected patient lymphoid cells then differentiated into cerebral organoids. After three months, the ALS-PDC affected organoids were smaller, their neurons had less extensive neurite outgrowth, and the organoids had more reactive astrocytes and M1 microglia, fewer resting and M2 microglia, and more open extracellular space. Most of these phenomena could be recapitulated by exposing unaffected organoids to *β*-methylamino L-alanine (BMAA), a toxic amino acid produced by cyanobacteria living with cycad plants. Furthermore, ALS-PDC affected organoids exhibited an exacerbated neuroinflammatory response to BMAA exposure via activation of caspase1/NLRP3 inflammasome. A genome-wide transcriptome analysis of the organoids showed that the most down-regulated pathways were taurine, alanine, aspartate, and glutamate metabolism; protein digestion; and absorption. The most down-regulated biological processes were type I interferon signaling, regulation of neuron differentiation and extracellular matrix organization. Our results suggested that the etiology of ALS-PDC is due to metabolic disorders that shifted microglia to a more pro-inflammatory M1 state instead of a non-inflammatory, repairing M2 state, which exacerbated inflammation and reduced extracellular matrix strength. Supplementation of transforming growth factor beta (TGF-β) to ALS-PDC affected organoids increased the expression of interferon-induced transmembrane proteins (IFITMs) and restored M2 microglia populations and extracellular matrix organization. Organoids containing networks of neurons, astrocytes, microglia derived from iPSC with our protocol provides an excellent cellular model for neurodegenerative disease modeling.

**Significance Statement:**

1. Western Pacific Amyotrophic Lateral Sclerosis and Parkinsonism–dementia Complex (ALS-PDC) cerebral organoids containing networks of neurons, astrocytes, and microglia were generated from patient specific lymphoid derived induced pluripotent stem cells.
2. ALS-PDC affected organoids were smaller, with neurons had less extensive neurite outgrowth, more reactive astrocytes and M1 microglia, fewer resting and M2 microglia, and more open extracellular matrix spaces when compared to ALS-PDC unaffected organoids.
3. Genome-wide transcriptome analysis indicated that ALS/PDC affected organoids had significantly lower expression of genes related to vitamin B6, amino acid and protein glycation metabolisms, down-regulated type I interferon signaling, the regulation of neuron differentiation and extracellular matrix production.
4. Our results implicated that the etiology of ALS-PDC is due to metabolic disorders that led the shift of microglia to more pro-inflammatory M1 state and less non-inflammatory resting, repairing M2 state of microglia subpopulation, which primed the exacerbated inflammation and reduced extracellular matrix strength. TGF-β promoted interferon-induced transmembrane protein (IFITMs) expression and restored the repairing M2 state of microglia population and extracellular matrix organization.

## INTRODUCTION

Western Pacific Amyotrophic Lateral Sclerosis and Parkinsonism-Dementia Complex (ALS-PDC) is a neurodegenerative disease that exhibits clinical and neuropathological similarities to Amyotrophic Lateral Sclerosis, Parkinson’s, and Alzheimer’s disease (Spencer et al., 2009, 2016; Miklossy et al., 2008). ALS-PDC was first identified among the Chamorro people of Guam. Studies showed that ALS-PDC in Guam was associated with chronic exposure to the traditional use of cycad seeds (*Cycads* spp.) for medicine and food among the Chamorro population (Zhang et al., 1996). One of the cycad toxins has been identified as **β**-*N*-methylamino-L-alanine (BMAA), which is produced by cyanobacteria that live on cycad plants (Kisby et al., 1992). BMAA affects the metabolisms of alanine, aspartate, and glutamate, as well as nitrogen metabolisms (Engskog et al., 2017; Delcourt, et al., 2018). To date, there is no convincing animal model for ALS-PDC. The explanation for this failure likely resides in differences in inflammatory response to metabolic changes between humans and animals. A comprehensive study revealed that gene dysregulation in mouse models of severe human inflammatory conditions did not correlate with human genomic changes (Seok et al., 2013). This highlights questions about the relevance of mouse models for studying the interaction between metabolic and immune systems, and their impact on neurodegenerative diseases.

Microglia are the resident immune cells of the central nervous system (CNS). They represent 10-15% of the total cell population in the CNS (Harry et al., 2013). Microglia can self-renew and their development depends on the IL34/CSF1R and TGF-β signaling pathways (Wang et al., 2012; Butovskyo et al., 2014). In response to damage-associated molecular pattern (DAMP) molecules such as ATP, reactive oxygen species (ROS), and advanced glycation end product (AGE) stimuli, microglia can be transformed into diverse activated states which have distinct cytokine secretion properties and phagocytic capacities through the activation of “sensome”, a cluster of proteins that include pattern-recognition receptors, purinergic receptors, cytokine and chemokine receptors, and extracellular matrix (ECM) receptors (Banati et al., 1993; Hickman et al., 2013, 2018). Studies showed that pro-inflammatory M1 microglia release pro-inflammatory cytokines, such as tumor necrosis factor-alpha (TNFα) and ROS, which promote NLRP3 mediated neuroinflammation and neuronal cell death (Sarlus et al., 2017; Tang et al., 2015). Pro-inflammatory M1 microglia mediated immune response was a key contributor to the pathogenesis of several neurodegenerative diseases including Alzheimer’s disease (AD) and Parkinson’s disease (PD) (Keren-Shaul, 2017; Moehle et al., 2015; Rodríguez-Gómez et al., 2020).

The non-inflammatory, repairing M2 state of microglia is primarily associated with promoting neurogenesis and extracellular composition reconstruction, and clearing aggregated proteins (Paolicelli, 2011; Mathys et al., 2017; Cowan, 2018). M2 microglia associated with secretion of anti-inflammatory and tissue repair cytokines such IL-4, IL-10, and TGF-β (Lana, et al., 2020). TGF-β can modulate the functional properties of microglia which plays a critical role in microglia homeostasis and prevent microglia from entering the neurotoxic M1 state (Brionne et al., 2003; Noh et al., 2016). In addition, TGF-β also plays an important role in maintaining the mechanical strength of connective tissues by increasing the expression of ECM proteoglycans and other fibrous matrix proteins (Diniz et al., 2012; Zoller et al., 2018; Kudo et al., 2018, Tzavlaki et al., 2020). However, the cellular mechanisms that govern the diverse phenotypes of microglia and their association of neurodegenerative diseases are not well understood. This limitation is due to the lack of cellular models that contain neurons, astrocyte, and microglia.

Stem cell technology has opened a new avenue for the role of microglia in neurodegenerative disease research (Yu et al., 2007; Fujimori et al., 2016, Dutta et al., 2017; Kelva et al., 2016; Yoon et al., 2019). Several methods for deriving microglia from human iPSCs have been developed (Muffat et al., 2016). However, most protocols differentiated iPSCs into mesodermal hematopoietic stem cells (EMPs), which were then directed to microglia. This study is the first report to derive patient-specific iPSCs from healthy and ALS-PDC patient lymphoid cell lines. Then, we described a robust method for generating cerebral organoids containing neuron-astrocyte-microglia from these iPSCs by using a chemically defined culture medium. Using cerebral organoids containing microglia, our results suggested that the imbalance of microglia population driven by metabolic disorders plays a key role in ALS-PDC disease progression. iPSC derived cerebral organoids containing neurons, astrocytes and microglia are an excellent cellular model to study the impact of genetic and environmental factors on microglia-mediated neurodegenerative diseases. They also present a platform for future studies targeting microglial phenotype switching for neurodegenerative disease treatment.

## RESULTS

### 1. Generation of cerebral organoids containing neuron-astrocyte-microglia networks from ALS-PDC affected and unaffected iPSCs derived from lymphoblast cells

Gender- and age-matched lymphoid cell lines (LCLs) were obtained from a patient with ALS-PDC and an unaffected individual (gifts from Drs. Teequ Siddique and Glen Kisby). As shown in Figure 1A, LCLs were cultured in RPMI 1640 complete media. After three days of culture, same cell density of LCLs (10,000/ml) were nucleofected with a mixture of episomal plasmids (pCEhOCT3/4, pCE-hSK, pCE-hUL, and pCE-mp53DD) that encode reprogramming factors (OCT3/4, SOX2, KLF4, LMYC, and LIN28) and mouse p53DD (p53 carboxy-terminal dominant-negative fragment) via neontransfection (Barrett et al. 2014). ALS-PDC unaffected iPSC colonies formed at day 25, while same size of ALS-PDC affected colonies appeared at day 32. The pluripotency of three cell lines of ALS-PDC affected and unaffected iPSCs were examined with stem cell factor OCT4 immunostaining (Figure 1A) and RNA-seq. RNA-seq results showed that both ALS-PDC affected and unaffected iPSCs expressed significantly higher levels of Nanog, Pou5F1, and Sox2 than that of LCLs (differentially expressed gene (DEG) read numbers 79, 37,1016 vs 63, 803,1016, respectively). Embryoid bodies (EB) which contain three germ layer cell lineages were derived from those iPSCs in a KnockOut serum medium (KSRM) supplemented with growth factors bFGF, EGF, LIF, and heparin under 10% CO_2_ culture conditions. Then EBs were further incubated with SB431542 and dorsomorphin for neuronal induction under 5% CO_2_ for 5 days (Aynun et al., 2015). Rosettes, neuronal tube-like structures containing neural progenitor cells (NPCs) and microglia progenitors (MPCs) were formed at day 20. Rosettes were immunostained with the neuroectodermal stem cell marker Nestin and microglia marker transmembrane protein 119 (TMEM119), the results showed that most microglia-like progenitors clustered at the center of rosettes. The number of microglia progenitors in the ALS-PDC affected rosette was significantly reduced when compared to controls (Figure 1B). Those rosettes were detached with STEMdiff Neural Rosette Selection Reagent and grown into organoids cultured in neuronal maintenance medium (NMM). After three months in culture, the average size of ALS-PDC affected organoids was 544 μm, which was only half the size of ALS-PDC unaffected organoids (1025μm) (Figure 1B; Supplemental Figure 1).

**Figure 1.**
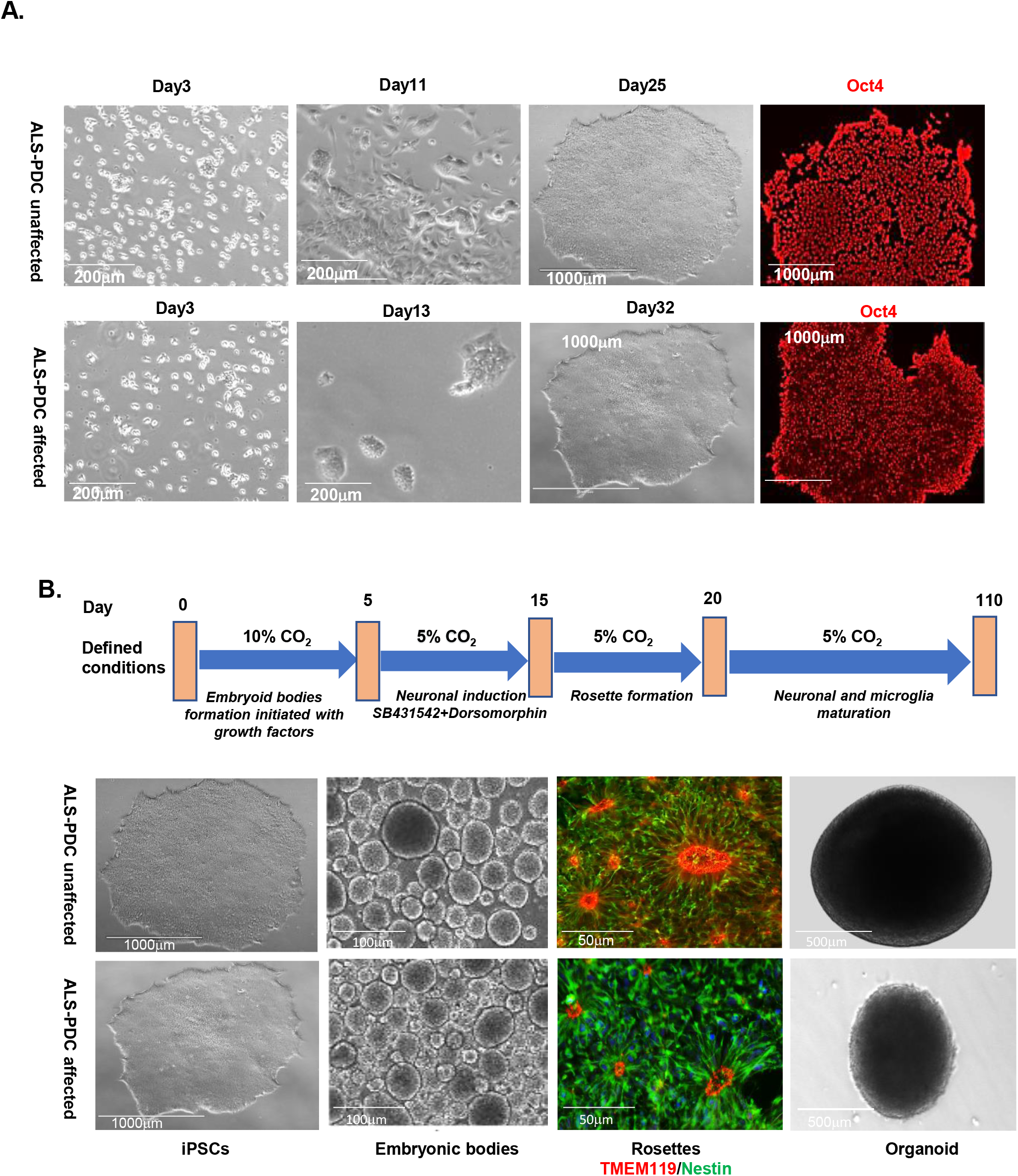
Generation of iPSCs and organoids from the LCLs of 57-year-old female Chamorro patient with ALS-PDC (*lower images*) and the age- and gender-matched 57-year-old healthy control subject (*upper images*). **A**. ALS-PDC unaffected LCLs at 3, 11, and 25 days and ALS-PDC LCLs at 3, 13, and 32 days following transfection with episomal plasmids. Representative iPSCs were immunostained with an OCT4 antibody. **B**. A schematic showing the procedure of derivation of cerebral organoids containing the neuron-astrocyte-microglia network. iPSCs were differentiated into embryoid bodies with growth factors under 10% CO_2_ culture condition, and then neuronal induction with SB 431542 and Dorsomorphin which converted them into rosette-forming neuroepithelial cells containing neuronal and microglia progenitors (stained by TMEM119/Nestin) under 5% CO_2_ culture condition. Rosettes were picked up to continue grow into 3D organoids. ALS-PDC affected organoids are smaller compared to control one (supplementary figure1).

### 2. ALS-PDC affected organoids had less extensive neurite outgrowth in neurons, more reactive astrocytes and M1 microglia, and less resting microglia phenotypes which exacerbated the inflammatory response to acute BMAA exposure

Three-month old ALS-PDC affected and unaffected cerebral organoids were treated with BMAA (100 μm) for two weeks. Untreated organoids were used as controls. These organoids were fixed and sectioned into 10-µm thick slices for immunostaining. Neurons and astrocytes were detected using Map2 (microtubule associated protein 2) and GFAP (glial fibrillary acidic protein). Resting microglia were detected using TMEM119 (transmembrane protein 119) and reactive microglia were detected using Iba1 (ionized calcium-binding adapter molecule). After three months in culture, ALS-PDC affected organoids had fewer developed neurons with less extensive neurite outgrowth and had distinct microglial and astrocyte phenotypes (Figure 2). In addition, ALS-PDC organoids had more open extracellular spaces, which caused some organoids to fall apart during culture. Morphology of the subpopulation of microglia was further examined in 2D cultures immuno-staining with TMEM119, Iba1 and M1 microglia marker CD86. The results showed that microglia derived from heathy control iPSCs displayed a more ramified morphology with elongated dendrites. Some of them changed to a brush shape in response to BMAA exposure. In contrast, microglia derived from ALS-PDC affected iPSCs were less ramified and had more diverse amoeboid shapes M1 positive microglia. In response to BMAA exposure, ALS-PDC affected microglia significantly increased M1, NLRP3 and caspase 1 expression (Figure 3A-l, 3B). Our results revealed that ALS-PDC affected organoids had more pro-inflammatory M1 microglia subpopulations which exacerbated the inflammatory response to acute BMAA exposure via caspase1/NLRP3 inflammasome activation.

**Figure 2.**
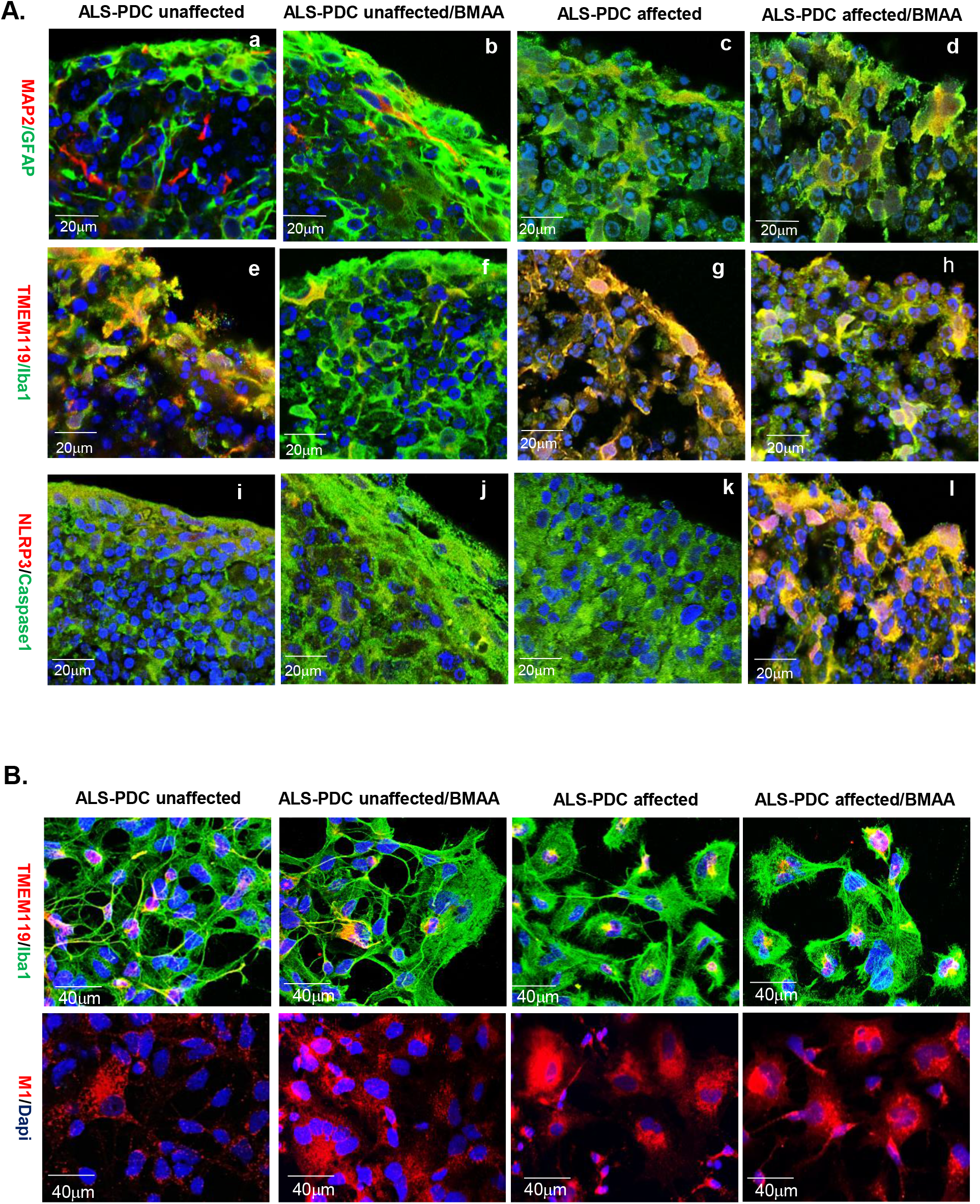
Comparison of neuron, astrocyte and microglia phonotype and their marker gene expressions both in 3D cerebral organoids and 2D culture from ALS-PDC-affected and unaffected iPSCs, along with BMAA treatments. **A**. Immuno-staining of tissues with different antibodies as listed on the left. Images were taken with Zeiss LSM 880 60X. Scale bars, 20 μm. **B**. The distinct morphologies microglia and M1 microglia population in both ALS-PDC affected and unaffected neuronal network were verified with TMEM119, Iba1 and M1 immuno-staining in 2D culture. Images were taken with confocal microscopy Zeiss LSM 880 20X Scale bar, 40 μm.

**Figure 3.**
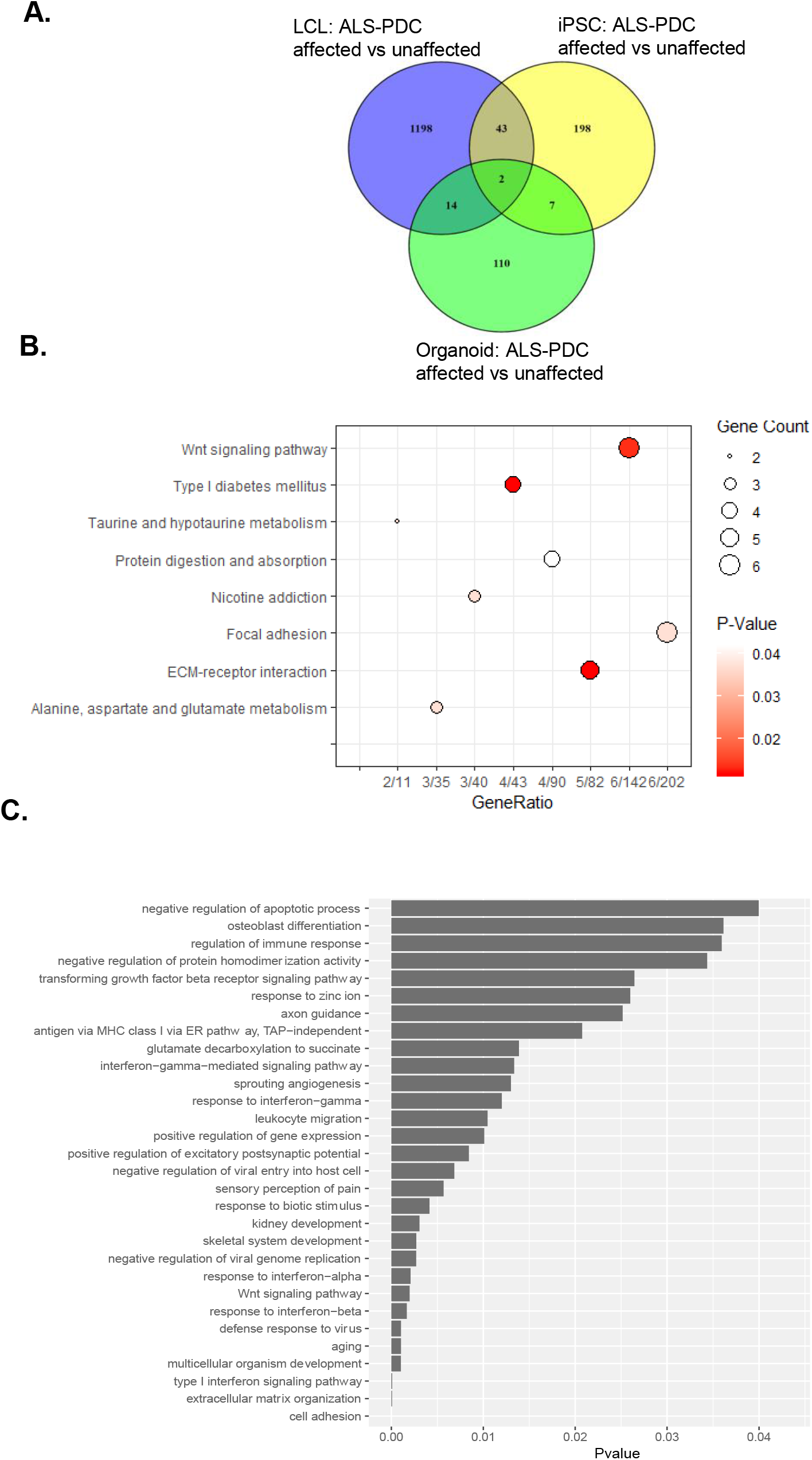
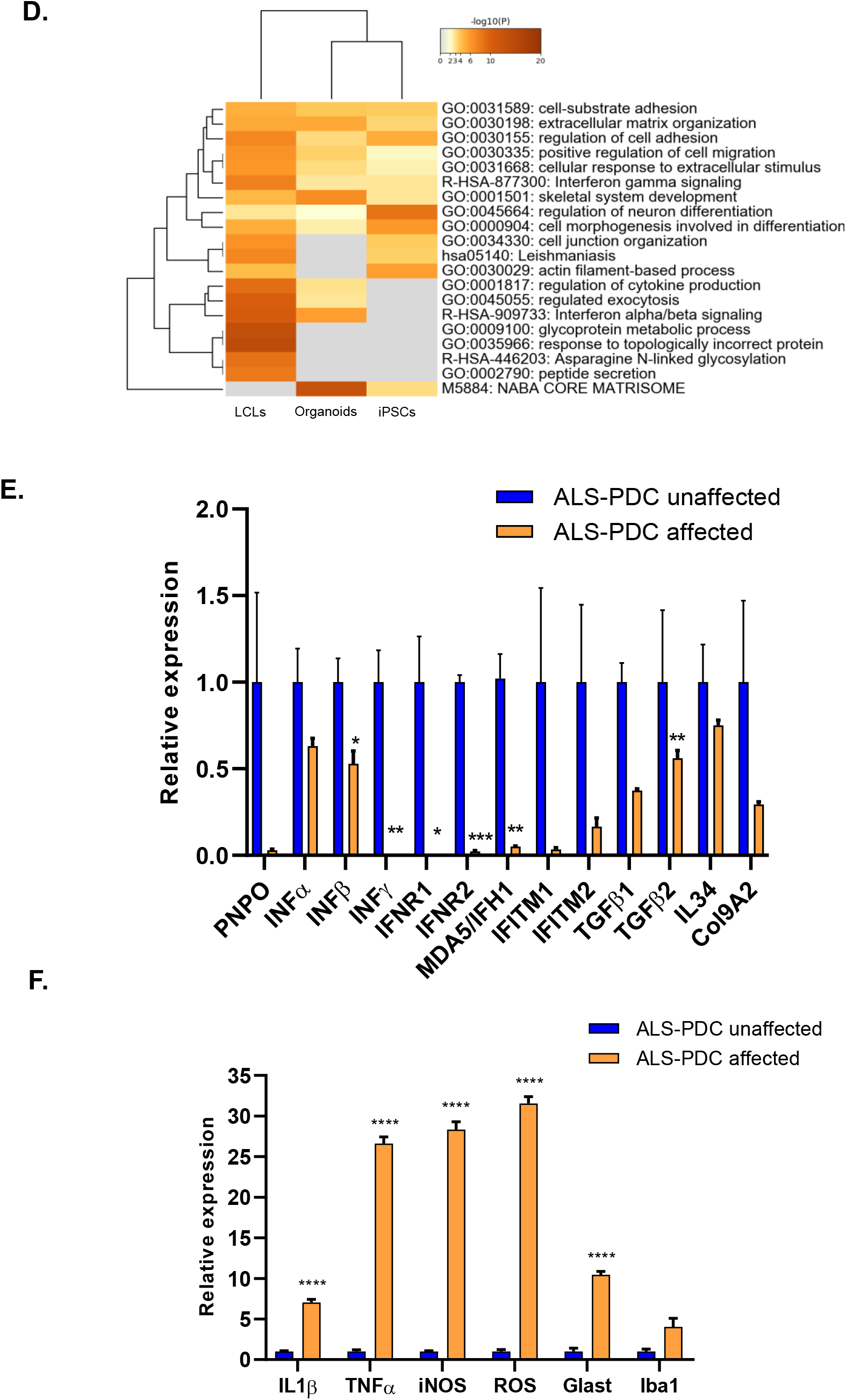
RNA-seq analysis by examining enriched transcripts through KEGG and GO analysis. **A**. The commonly down-regulated genes between LCL, iPSCs and organoids. **B**. Significant down regulated KEGG pathways compared between ALS-PDC affected and unaffected organoids. **C**. GO term enrichment analysis down biological process ALS-PDC affected vs unaffected organoid. **D**. The molecular signatures of GO term enrichment associated LCL, iPSC, and cerebral organoids across the different cell types. **E & F**. RT-qPCR confirmed the selected genes expression. The data were analyzed via Student’s t-test by software Graph Pad Prism 8. All values are compared to ALS-PDC unaffected and expressed as mean±SEM. ****p<0.0001 **p<0.001

### 3. Transcriptome analysis revealed that ALS/PDC affected organoids had significantly lower expression of genes related to metabolism, immune response, cytoskeletal reorganization, and neurogenesis

To further understand the molecular mechanism of ALS-PDC etiology, a transcriptome analysis was performed. To this end, RNA was isolated from ALS-PDC affected and unaffected LCLs, iPSCs, and cortical organoids. RNA-seq analysis was performed to identify differentially expressed genes (DEG, defined as 2-fold up or down regulated with *p*-value <0.05) in ALS-PDC affected cells compared to unaffected ones. At the LCL stage, there were 1373 up-regulated and 2083 down-regulated DEGs; after reprogramming LCLs into iPSCs, 460 genes were up-regulated and 560 down-regulated. At the organoid stage, 4 genes were up-regulated and 133 down-regulated (Supplemental table 1). Among them 14 commonly down-regulated gene clusters between LCL and organoids were immunity related proteins (IFITM 1&3, OAS3, BTN3A2), glutamate receptors and neurogenesis proteins (GRM8, NRGN), and cytoskeletal reorganization and cell-adhesion proteins (Col9A2, ITGA4) (Figure 3A). KEGG pathway analysis of LCL and organoid DEGs showed that at the LCL stage, the most down-regulated genes are in metabolic pathways (Supplemental figure 2). At the organoid stage, the most down-regulated pathways are taurine, alanine, aspartate, and glutamate metabolisms, protein digestion and absorption, ECM-receptor interaction, and immune response (Figure 3B). GO term enrichment analysis of ALS-PDC affected vs unaffected organoids indicated that the most significant down-regulation of the biological process were cell adhesion, extracellular matrix organization, and the type I interferon signaling pathway (Figure 3C). Those GO term enrichment signatures were consistent across LCL, iPSC, and cerebral organoids different cell types (Figure 3D). Selected gene expressions that were particularly focused on the metabolic enzymes and type I interferon signaling pathway were validated with reverse transcription and real-time PCR (RT-qPCR). The results indicated that ALS-PDC affected organoids expressed very low levels of Pyridoxine/pyridoxine 5’-phosphate oxidase (PNPO), interferon gamma, interferon production regulators, interferon-induced membrane proteins (IFITM), interferon induced helicase C domain 1MDA5/IFH1, Interleukin-34 (IL34), TGF-β, and Col9A2 (Figure 4E) while significantly up-graduating the expression of IL1β, TNFα, iNOS, ROS glutamate aspartate transporter (GLAST), and Iba1 (Figure 4F).

**Figure 4.**
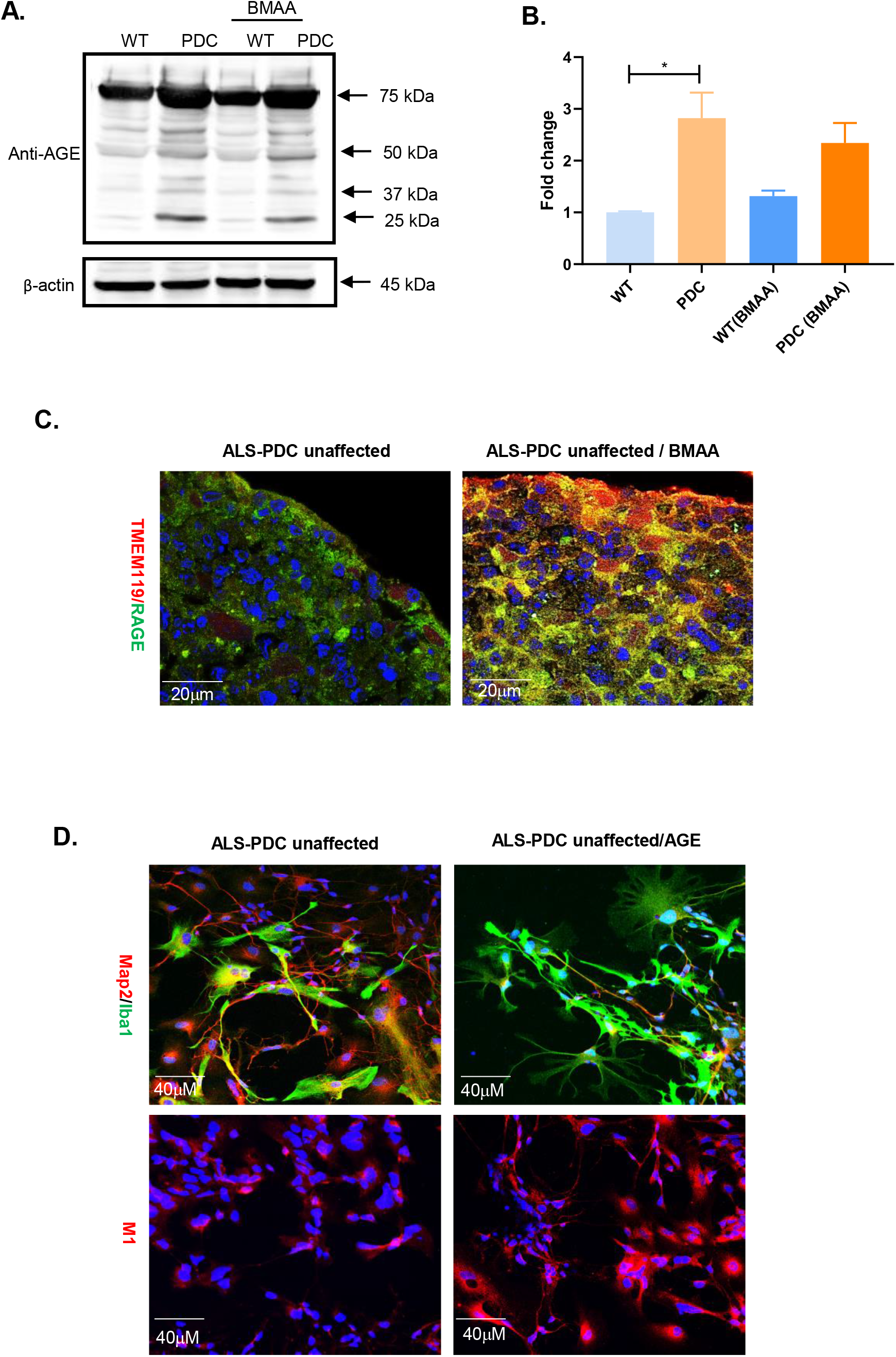
ALS-PDC affected organoids accumulated high level of advanced glycation end products that lead microglia activation. **A**. Western blotting indicated higher level of AGE protein modification in ALS-PDC unaffected organoids. β-actin is a loading control. **B**. Quantification of major protein modification bands. All values are compared to ALS-PDC unaffected and expressed as mean±SEM. *p<0.01. **C**. Cycad toxin BMAA exposure increase advanced glycation end products receptor RAGE expression and promoted microglia activation. **D**. AGE mediated microglia activation were verified in 2D culture. Images were taken with confocal microscopy Zeiss LSM 880 20X Scale bar, 40 μm.

### 4. ALS-PDC affected organoids accumulated high levels of advanced glycation end products that promote microglia activation

PNPO is a critical enzyme for the production of pyridoxal 5’-phosphate (PLP), the active form of vitamin B6. Vitamin B6 regulates the balance of amino acids, protein metabolism, and the production of antioxidants by counteracting the formation of ROS and advanced glycation end-production (AGEs) (Feng et al., 2018). AGEs are heterogeneous groups of irreversible adducts formed by the glycation of proteins with reducing sugars (Prasad et al., 2018; 2019). Western blotting confirmed that ALS-PDC affected organoids accumulated a much higher level of AGE modified proteins compared with unaffected ones (Figure 4 A&B). Furthermore, immunostaining of ALS-PDC unaffected organoids for AGE receptors (RAGE) and TMEM119 indicated that BMAA exposure promoted increased RAGE expression and microglia activation. Combine with iNOS RT-PCR result suggested that the AGE mediated microglia activation via AGE/RAGE/iNOS pathway which pathway had been shown to be involved in inflammation (Prasad et al., 2018 & 2019; Younessi, et al., 2011). In addition, AGE mediated microglia activation was further verified in 2D neuronal network. As showed in figure 4D. The results indicated that AGE exposure significantly altered microglia morphology and increased M1 microglia population. These results supported our hypothesis that the etiology of ALS-PDC is due to metabolic disorders that polarized microglia into more pro-inflammatory M1 state and promoted neuroinflammation.

### 5. ALS-PDC affected organoids reduced the M2 microglia population. TGF-β promoted interferon response protein IFITM expression and restored the M2 microglia population and extracellular matrix organization

Our RNA-seq data indicated that ALS-PDC affected organoids had significantly lower expression of genes related to the type I interferon signaling pathway. Interferon response protein IFITM had been shown to be involved in microglia polarization (Wee et al., 2015). ALS-PDC affected and unaffected organoids were further examined with immunostaining of IFITM with M1 microglia marker CD86 (Differentiation 86) and M2 microglia marker CD206 (mannose receptor). The result showed that IFITM co-expressed with CD206 in M2 microglia but not in M1 microglia. Furthermore ALS-PDC affected organoids had fewer M2 microglia (Figure 5 A&D). BMAA exposure increased the IFITM expressed microglia in ALS-PDC unaffected organoids but failed to do so in ALS-PDC affected organoids indicating that the type I interferon signaling pathway deficiency in ALS-PDC affected organoids plays an important role in M2 microglia polarization (Figure 5A). In addition, cytokine secretion tests for IL1β, TNFα, IL10, IL6, IL4, and TGF-β showed that ALS-PDC affected cultures secreted 60% less TGF-β (Figure 5C) compared with the healthy control. We observed less developed neurons in ALS-PDC affected organoids. We hypothesize that M2 microglia deficiency could be due to the deficiency of interleukin 34 or TGF-β. Supplementation of interleukin 34 (100 ng/ml) and TGF-β (50 ng/ml) cytokines to ALS-PDC affected organoids showed that TGF-β increased IFITM expression and M2 microglia population significantly (Figure 5D&E). In addition, TGF-β restored cell-cell connections and increased ECM production by promoting the distribution of periostin, a secretory protein that binds to type I collagen and fibronectin and contributes to the increased mechanical strength of connective tissues (Figure 5 D&E).

**Figure 5.**
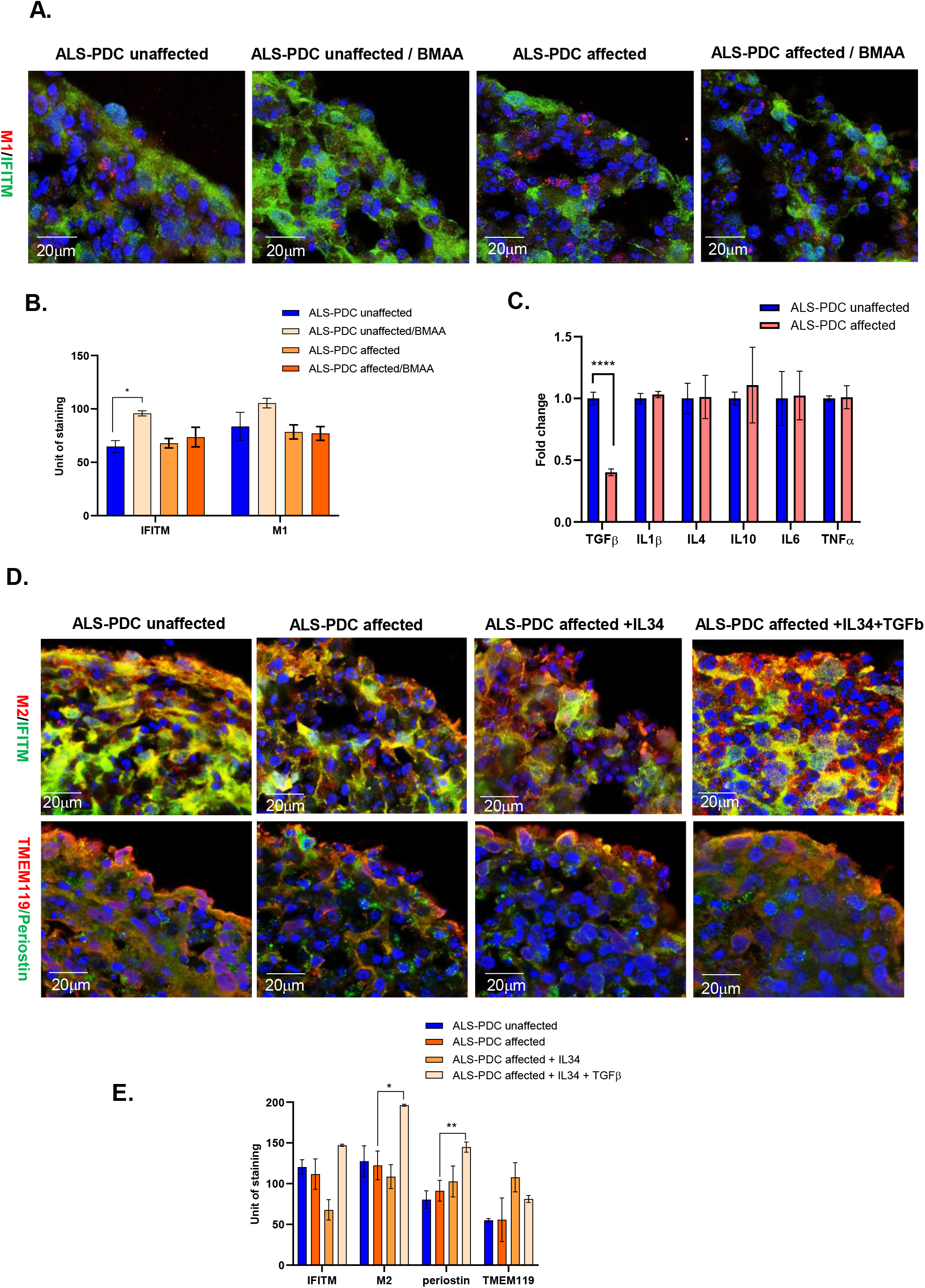
ALS-PDC affected organoids reduced repairing M2 microglia population. TGF-β restored M2 microglia and ECM strength in ALS-PDC affected organoids. **A**. BMAA treatment increased the expression of IFITM in ALS-PDC unaffected organoids, but not ALS-PDC affected one. Scale bar, 20 μm **B**. Analysis of fluorescence intensity of IFITM, M1 and M2. **C**. Cytokine secretion tested with IL1β, TNFα, IL10, IL6, IL4 and TGF-β. **** p<0.0001. **D**. TGF-β treatment increased the expression of IFITM and promoted M2 microglia polarization, periostin redistribution and restored ECM strength of ALS-PDC affected organoids. **E**. Analysis of fluorescence of IFITM, M2, TMEM119, Periostin in ALS-PDC unaffected and affected organoids treated with IL34 and TGF-b. All the data compared to ALS PDC unaffected **** p<0.0001, ***p<0.001,**p<0.01. Error bars=SEM

In summary, our results concluded that the etiology of ALS-PDC is due to metabolic disorders that led the shift of microglia to more pro-inflammatory M1 state and less non-inflammatory resting, repairing M2 state of microglia subpopulation. Imbalance of microglia population leads to an exacerbated inflammatory response and reducing extracellular matrix production in ASL-PDC affected organoids and thus increase neuronal cell death. Furthermore, our result suggested that TGF-β can restore M2 microglial homeostasis and enhance the strength of the extracellular matrix. Our directed differentiation protocol provided an excellent platform to further characterize the biological roles of microglia and their interactions with neurons, astrocytes, and extracellular matrices, both in healthy and pathophysiological conditions.

## DISCUSSION

Metabolic and immune systems are the most fundamental requirements for survival (Paludan et al., 2020). Microglia is a key player to integrate the metabolic regulation and immune response. Microglia homeostasis and proper functions is critical to CNS health. Experimental studies demonstrated that microglia display an enormous plasticity in their response to environmental insults. Activated microglia participate in defense, phagocytosis, uptake and clearance of Aβ, and tissue regeneration. Sustained chronic inflammatory states due to nutritional imbalances or neurotoxin containing foods can lead to change microglia homeostasis and elevates risk for microglia associated disorders (Khoury et al., 2010). Despite the critical role of microglia in brain physiology and pathologies, currently little is known about the cellular mechanisms involved in shifting microglial populations and phenotypes and their impact on neurodegenerative diseases due to the lack of a cellular model that contains a diversity of multiple cell types and ECM. In this study, for the first time, control and ALS-PDC patients’ lymphoblast cells were utilized to generate patient-specific iPSCs. From these iPSCs, cerebral organoids containing neurons, astrocytes, microglia, and ECM were created using a directed differentiation approach to model the etiological role of cycad toxin BMAA in ALS-PDC. Our results indicated that ALS-PDC affected organoids had fewer developed neurons, more reactive astrocytes and M1 microglia, fewer resting and M2 microglia, and more open ECM spaces when compared to ALS-PDC unaffected organoids.

Transcriptome analysis suggests that the genes most significantly affected by the cycad toxin BMAA are in metabolic pathways, the regulation of neuron development, immune response, and extracellular matrix organization which are preserved across the different cell types. The results from RT-qPCR, western blotting, and immunostaining confirmed that the cycad toxin BMAA increased AGE/RAGE/iNOS expression which promote M1 microglia activation. Furthermore, we also observed that ALS-PDC organoid has less M2 microglia population. We examined the involvement of the defective interferon signaling pathway and interferon induced protein IFITM in repairing M2 microglia polarization. Our study suggested that interferon signaling played a significant role in M2 microglia polarization. Deficiency of TFITM altered microglia morphology and properties so that they were unable to interact with neurons and astrocytes or clear debris, thereby leading to toxic microenvironments. Although the importance of interferons in the antiviral response has long been appreciated, the relationship between interferons, metabolic disorder and microglia polarization needs to be further investigated (Wee et al., 2015). Future studies can employ gene knock out and single-cell gene expression methodology to identify interferon mediated molecular mechanisms underlying the phenotypic switch and driving microglia toward beneficial functions.

Microglial proliferation and polarization are highly dependent on the constant communication with astrocytes and neurons through receptors and cytokines. Their interactions also function as a feedback loop to maintain microglia homeostasis and prevent M1 microglia reaction (Norden *et al*, 2015b, 2016). We have identified TGF-β as a major differentiation factor for microglia maturation, polarization, and remodeling of the ECM structure. The results showed that supplementation with TGF-β can increase the repairing M2 microglia population. Our differentiation platform will further address the interactions among microglia, astrocytes, neurons, and the extracellular matrix in our future study.

In summary, we have developed a platform to obtain patient derived organoids containing neurons, astrocytes, microglia, and extracellular matrices for disease modeling. This system recapitulates normal human brain development and neurodegeneration in response to neurotoxin BMAA exposure. It provides a cellular model for new insights into the role of microglia in neurodegenerative disease due to genetic variation or the impact of environmental factors. Furthermore, microglia could represent a promising pharmacological agent for future neurodegenerative disease therapy.

## EXPERIMENTAL PROCEDURES

### 1. Generation of patient-specific iPSCs from ALS-PDC affected lymphoid cell lines

Gender- and age-matched ALS-PDC affected and unaffected lymphoid cell lines cells (LCLs) were gifts from Drs. Teequ Siddique and Glen Kisby. LCLs cells were cultured in RPMI 1640 complete media until confluence. The patient-specific iPSCs were generated from LCLs by transfection with a combination of episomal plasmids (pCE-hOCT3/4, pCE-hSK, pCE-hUL, and pCE-mp53DD) (Addgene Inc.), as previously reported (Barrett et al. 2014). Approximately, a 2.5 µg equal amount mixture of pCE-hOCT3/4, pCE-hSK, pCE-hUL, and pCE-mp53DD were co-transfected into ∼ 1.0×10^6 LCLs via neontransfection (NEON transfection). The transfected LCLs were then plated into a matrigel-coated six-well plate and maintained in TeSr-E7 xeno-free containing media (Stem Cell Tech Inc.). Twelve days after transfection, the TeSr-E7 media was slowly replaced with mTeSR-1 media. Around 20-30 days after transfection, colonies that exhibited morphologies resembling human embryonic stem cells were picked and plated onto matrigel-coated six-well plated in mTesr-1 media. Soon after, the putative iPSC colonies were replaced onto matrigel-coated T25 flasks in mTeSR-1 media. The iPSC colonies were grown in matrigel-coated T25 flasks with mTeSR-1 media and passaged every 4 days. Pluripotency was determined by examining the iPSCs for OCT4 expression by immunostaining.

### 2. Differentiation of organoids containing neurons-astrocytes-microglia network from ALS-PDC affected and unaffected iPSCs

iPSCs from ALS/PDC affected and unaffected patients were expanded and enzymatically dissociated with collagenase IV (2 mg/mL, Life Technologies Inc.). Dissociated cells were transferred into a low adhesion suspension culture plate (Olympus, Inc.) cultured in Knock out serum replacer (KSRM) (Life Technologies Inc.) supplemented with 10 ng/ml bFGF (Life Technologies Inc.), 10 μM ROCK inhibitor (Tocris Bioscience Inc.), 50 ng/ml EGF (R&D Systems Inc.), 1000 Unit/mL LIF (Millipore Inc.) and 1 μg/mL heparin (Sigma-Aldrich Inc.) incubated at 37°C under 10% CO_2_ culture conditions for 5 days to derive a three germ layer embryonic bodies (EBs). The EBs were further cultured in neuronal induction medium (NIM) which is a 1:1 combination of SKSRM and neuronal maintenance medium (NMM) supplemented with 10 μM SB431542 and 1 μM dorsomorphin (both from Tocris Bioscience Inc) for 10 days for neuronal induction. NMM is a 1:1 mixture of DMEM/F12 and Neurobasal™-AMedium (both from Gibco Inc.) supplemented with N-2 Supplement and B-27™ Supplement (Gibco Inc.), as well as 5 µg/mL insulin (Tocris Bioscience Inc), 1 mM GlutaMAX, 100 µM nonessential amino acids and 100 μM β-mercaptoethanol (Gibco Inc.). Over the 10 days incubation, SKSRM media was gradually replaced with NMM media by increasing the ratio of NMM versus SKSRM 25% every two days. The media shifted from 100% SKSRM to 100% NMM with 5 media changes. After neuronal induction, the neurospheres containing neuronal and microglia progenitors were collected at 400 g for 5 min, and then were gently suspended with gentle cell dissociation solution (Stem Cell Technologies Inc.) and incubated for 7 min at 37°C, and then broken down into smaller fragments by using a 5 ml polystyrene serological pipette. The cells will be placed to matrigel-coated plates and culture in NMM under 37°C, 5% CO2 culture condition. After 3-5 days, neuroepithelial sheets formed, which contained neural rosettes with neuronal and microglia progenitors. For the generation of organoids, the neural rosettes were detached with incubation of STEMdiff™ Neural Rosette Selection Reagent (Stemcell Technologies) for 30 min at 37°C. Individual rosettes were transferred into separate wells of a low adhesion suspension 24 well culture plate. The organoids were maintained with NMM with or without supplement of interleukin 34 (100 ng/ml), CSF (5 ng/ml), and TGF-β (50 ng/ml).

For the generation of a monolayer neuronal culture, neuronal rosettes were treated with Gentle Dissociation Reagent (Stemcell Technologies Inc.) at 37°C for 10 min and placed at a density of approximately 1×10^4^ cells in matrigel-coated plates with coverslips. After culture in NMM supplemented with interleukin 34 (100 ng/ml), CSF (5 ng/ml), and TGF-β (50 ng/ml) for two weeks, the progenitors grew into mature neurons, astrocytes and microglia.

### 3. Cycad toxin BMAA exposure

For BMAA dosing study, neuronal rosettes were dissociated and seeded in 24-well plates at a density of 50,000 cells/cm^2^ and culture in NMM for 2 weeks. Mature neuronal network were exposed to varying concentrations of 1 µM, 10 µM, 100 µM, and 1000 µM BMAA ordered from Sigma (Cat# B-107) for 3 days. 100 µM of BMAA was selected to expose to the patient-specific organoids for 14 days. The neurotoxins were then reintroduced into the wells every time the culture media was replaced.

### 4. Organoid processing and immunohistochemistry staining

Organoids were fixed in 4% PFA for 24 hours and then placed into 30% sucrose solutions at room temperature. Before sectioning, organoids were transferred into molding cup trays (Polysciences Inc.) filled with OCT compound (Tissue Tek Inc.) and flash frozen using liquid nitrogen. The frozen OCT block containing the organoid was then sectioned (10 µM thick) using a cryostat (Leica, Model CM 1950, Germany). For the immunocytochemistry staining, the slices were washed with PBS three times and then permeabilized in 0.1% Triton X-100 (Sigma-Aldrich) for 15 min at room temperature. After the permeabilization, the slices were blocked with 2% BSA, and incubated overnight with primary antibodies markers of mature neurons and astrocyte Map2 (Santa Cruz, #SC20172)/GFAP (ThermoFisher #MA5-12023), microglia markers: TMEM119 (Abcam 185333), Iba1 (Abcam #15690, IFITM (sc-374026), CD86 (ab239075), CD208 (ab8918), Inflammasome markers NLRP3 (PA5-79740), Capase 1(Santa Cruz, SC-392736), Periostin/ (SC-398631) at 4°C in a humidity chamber to prevent evaporation. The next day, the slices were washed two times with washing buffer (1×PBS containing 0.1% Tween 20) and twice with PBS), then the slices were incubated for 2 hours at room temperature with fluorescent secondary antibodies (Life Technologies Inc.). After 2 hours of incubation, the organoid slices were washed three times with washing buffer and once with PBS. Finally, the organoid slices were cover-slipped with Fluoromount-G mounting medium containing DAPI. Immunohistochemistry images were taken with Zeiss LSM 880 with Airyscan confocal microscope. Data were collected from three independent images and were analyzed with Image J software (https://imagej.nih.gov/ij/) and analyzed via ANOVA and Student’s t-test. P < 0.05 was considered statistically significant. Correlations will be determined by calculating Pearson coefficients.

### 5. RNA-seq and data analysis

LCLs, iPSCs and organoids were harvested and lysed with TRIzol™ Reagent (Invitrogen Inc.) and total RNA was extracted according to the manufacturer’s instructions. RNA-seq library construction, sequencing, and analysis were performed at the City of Hope Integrative Genomics Core following routine as described. Libraries were prepared with Kapa RNA mRNA HyperPrep kit (Kapa Biosystems Inc.) according to the manufacturer’s protocol. Reads were aligned against the human genome (hg19) using TopHat2 (Kim et al., 2013). Read counts were quantified using htseq-count (Anders et al., 2015), with UCSC known gene annotations (TxDb.Hsapiens.UCSC. hg19.knownGene (Hsu et al., 2006). Fold-change values were calculated from Fragments Per Kilobase per Million reads (Mortazavi et al.,2008) normalized expression values, which were also used for visualization (following a log_2_ transformation). Aligned reads were counted using GenomicRanges (Lawrence et al., 2013). Prior to p-value calculation, genes were filtered to only include transcripts with an FPKM expression level of 0.1 (after a rounded log2-transformation) in at least 50% of samples (Warden et al., 2013) as well as genes that are greater than 150 bp. The general scripts are a modified version of a template for RNA-Seq gene expression analysis (https://github.com/cwarden45/RNAseq_templates/tree/master/TopHat_Workflow).RNA-Seq data for this study is publicly deposited in NCBI with accession number X (in submission; the accession number will be provided soon).

For the ALS/PDC affected vs unaffected comparison (n=3), p-values were calculated from raw counts using DESeq2 (Love et al., 2014), and the false discovery rate (FDR) values were calculated using the method of Benjamini and Hochberg (Benjamini et al.,1995). Differentially expressed genes were defined as those with fold-change >2.0, unadjusted p-value < 0.05, and FDR < 0.25. Gene Ontology enrichment was calculated using goseq (Ashburner et al., 2000). Gene symbol GO mappings were defined within goseq for the genome reference “hg19”. A histogram was created from the results using plotting -log10 (p-value) for gene (Young et al., 2010). Selected genes were validated with reverse transcription and real-time PCR (RT-qPCR) with SYBR Premix on Bio-Rad CFX 96 (Bio-Rad Inc.), according to the manufacturer’s instructions. All primer sequences are shown in Table S2. The rich factor is the ratio of differentially expressed gene numbers of the pathway to all gene numbers annotated in this pathway.

### 6. Cytokine secretion assay

Cytokine secretion assays were performed using Human Multi-Analyte ELISArray plate (Qiagen CMEH6321A). IL1β, TNFα, IL10, IL6, IL4 and TGFβ Antigen Standard Cocktail were prepared and used as positive controls following the manufacturer’s recommendation. The ALS-PDC affected and unaffected cell culture supernatants were collected and centrifuged for 10 min at 1000x g. 50 µl of each sample and Antigen Standard Cocktail positive control samples were transferred into each well of an ELISArray plate containing 50 µl assay buffer. The plate was gently tapped for 10 seconds and incubated for 2 h at room temperature. After washing the ELISA well with 350 µl 1X washing buffer three times, 100 µl Detection Antibody was added and incubated for 1 hour at room temperature. 100 µl Avidin–HRP solution was added after three washes with washing buffer to each well and the plate was incubated for 30 min at room temperature. The ELISA wells were washed for 4 times and 100 µl Development Solution was added to each well. The plate was incubated for 15 min at room temperature in the dark, after which 100 µl of Stop solution was added to each well. The color changes from blue to yellow were read by OD 450 nm within 30 min of stopping the reaction. All values are expressed as mean±SD and the software GraphPad Prism 8 was used. Comparison of means was conducted using t-test and results were considered statistically significant if p-value was < 0.05.

### 7. Western blot analysis

ALS-PDC affected and unaffected cells along with BMAA treated cells were harvested and homogenized in cold RIPA buffer (0.15 M NaCl, 1% NP-40, 0.05% deoxycholic acid, 1% SDS, and 50 mM Tris) containing protease inhibitors (Roche Inc.). Equal amounts of protein (50 μg) were resolved by 10% SDS-PAGE and then transferred onto polyvinylidene difluoride membranes. Membranes were probed with specific antibodies with AGE (Abcam # 23722). β-actin (Cell Signaling Technology, #4970) was used as control. Data were collected from at least three independent tests, and the results were expressed as mean ± SEM. Statistical analyses were performed with SPSS 19.0 software (IBM, Armonk, NY, USA). Multiple group comparison was conducted with one-way ANOVA, which followed by Tukey post hoc test. Values of *p* < 0.05 were considered as statistically significant.

## FIGURE LEGENDS

**Figure S1. Organoid size comparisons. A**. Phase contract images of selected image of ALS-PDC affected and unaffected organoids. **B**. Quantification the size of organoids. All the data from three separate images compared to ALS-PDC unaffected **p<0.01. Error bars=SEM.

**Figure S2**. Significant down regulated KEGG pathways compared between ALS-PDC affected and unaffected LCL.

**Table S1**. The number of differentially expressed Genes.

**Table S2:** Primers used for q-PCR analysis

## ACKNOWLEDGEMENT

We would like to thank Drs. Teequ Siddique and Glen Kisby for the LCL cell lines, City of Hope Integrative Genomics Core providing the support of RNA-seq and data analysis and the support of Western University of Sciences through an intramural grant.

## Author’s contributions

Y. Hong: conceived and designed the project; data analysis and manuscript writing. X. Dong: generation of iPSC cell lines and perform cell culture, RNA isolation, RT-qPCR and immunohistochemistry staining. L. Change: development of the procedure of generation of organoids from iPSCs and performed immunohistochemistry staining. M. Chang: perform cell culture and immunohistochemistry staining. C. Xie: perform cytokine secretion and western blotting analysis. J. S. Aguilar: Perform q-PCR and data analysis. Q.Q. Li: co-mentor students, data analysis and manuscript editing.

## Graphic Figure

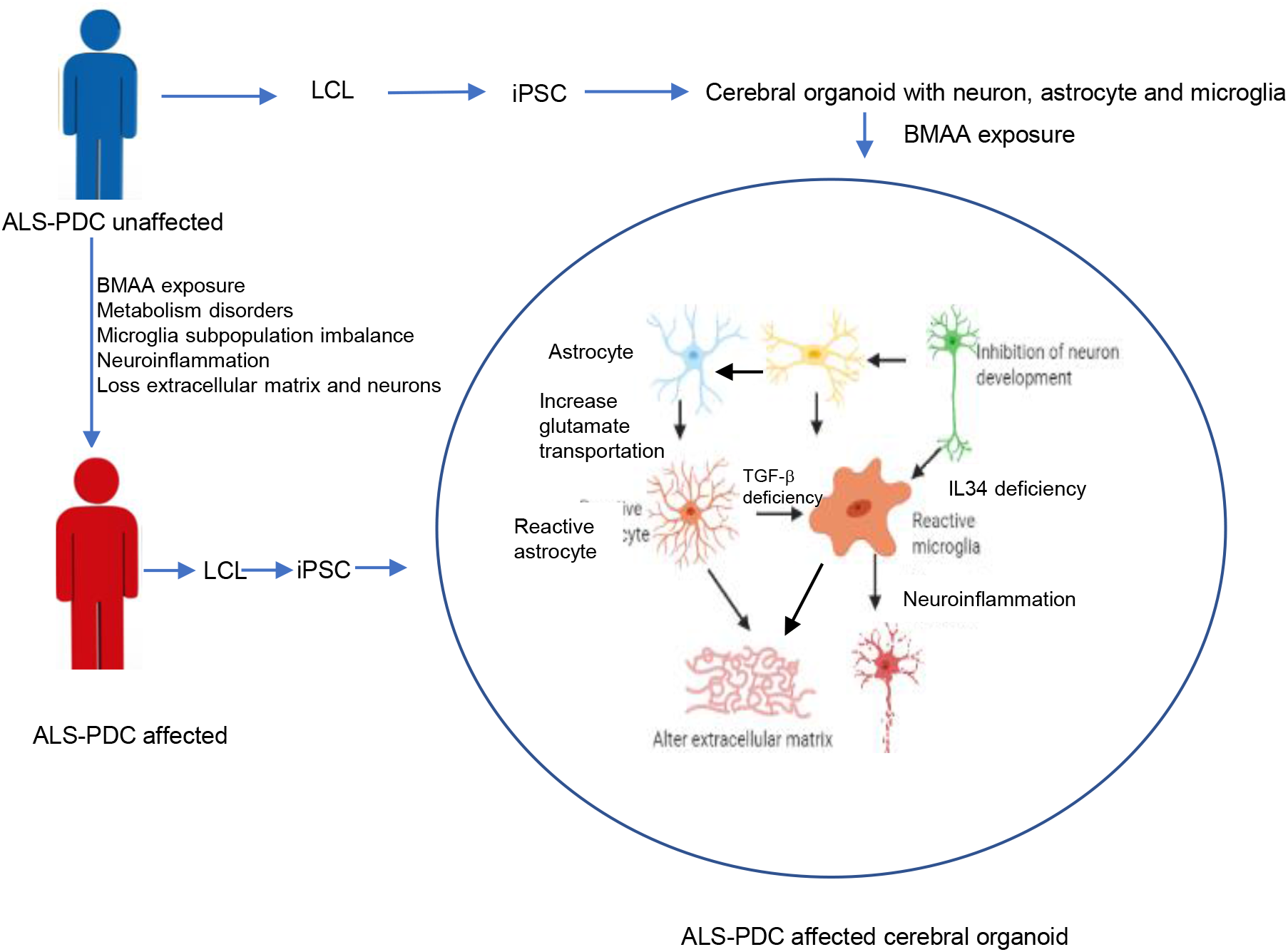

## Supplementary Figures

**Figure S1.**
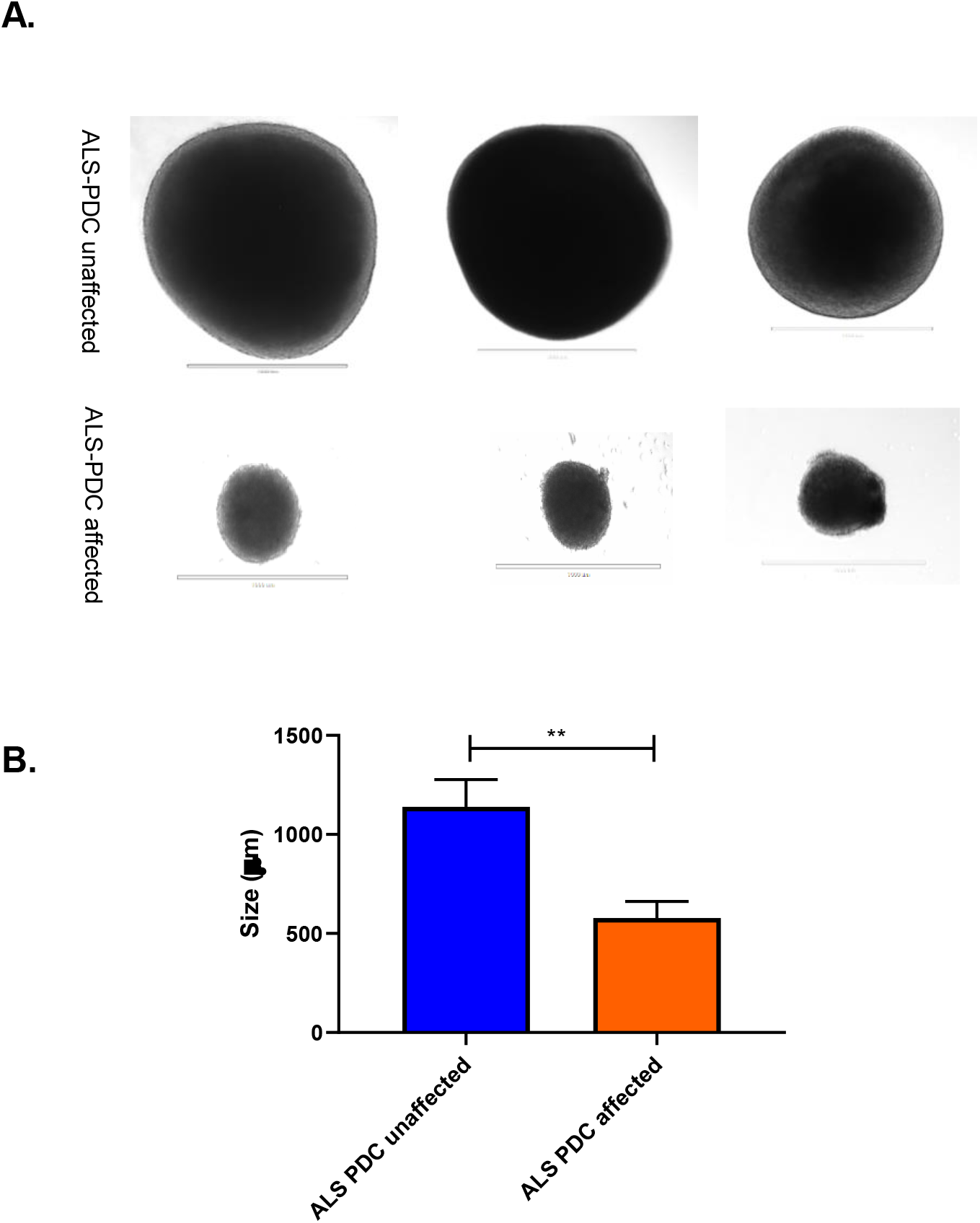
Organoid size comparisons. **A**. Phase contract images of selected image of ALS-PDC affected and unaffected organoids. **B**. Quantification the size of organoids. All the data from three separate images compared to ALS-PDC unaffected **p<0.01. Error bars=SEM.

**Table S1.** The number of differentially expressed Genes.

| Comparison | Up-regulated | Down-regulated |
| --- | --- | --- |
| LCL: ALS-PDC affected vs unaffected | 1,373 | 2,083 |
| iPSC: ALS-PDC affected vs unaffected | 460 | 560 |
| Organoid: ALS-PDC affected vs unaffected | 4 | 133 |

**Figure S2.**
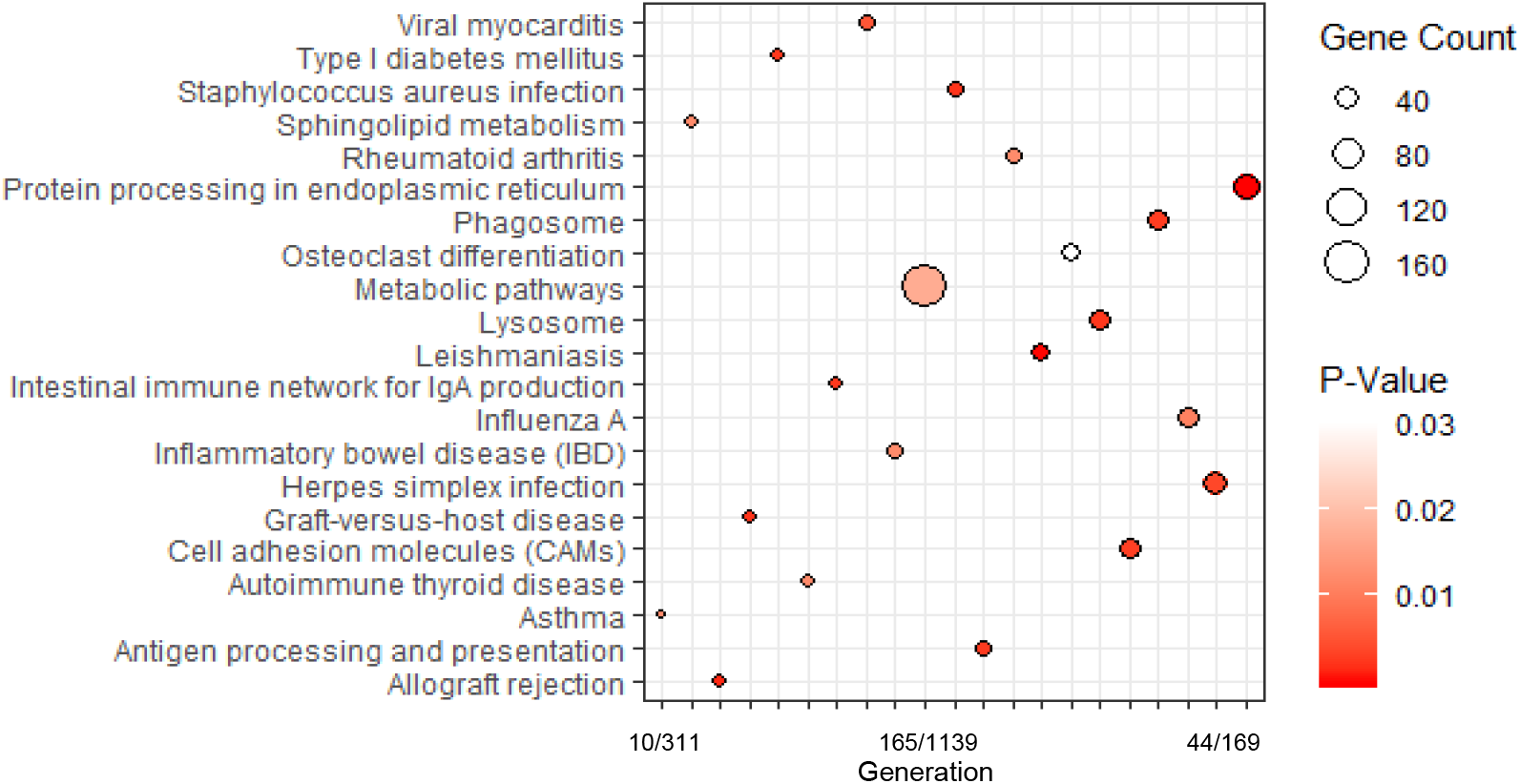
Significant down regulated KEGG pathways compared between affected and unaffected LCL.

**Table S2.**
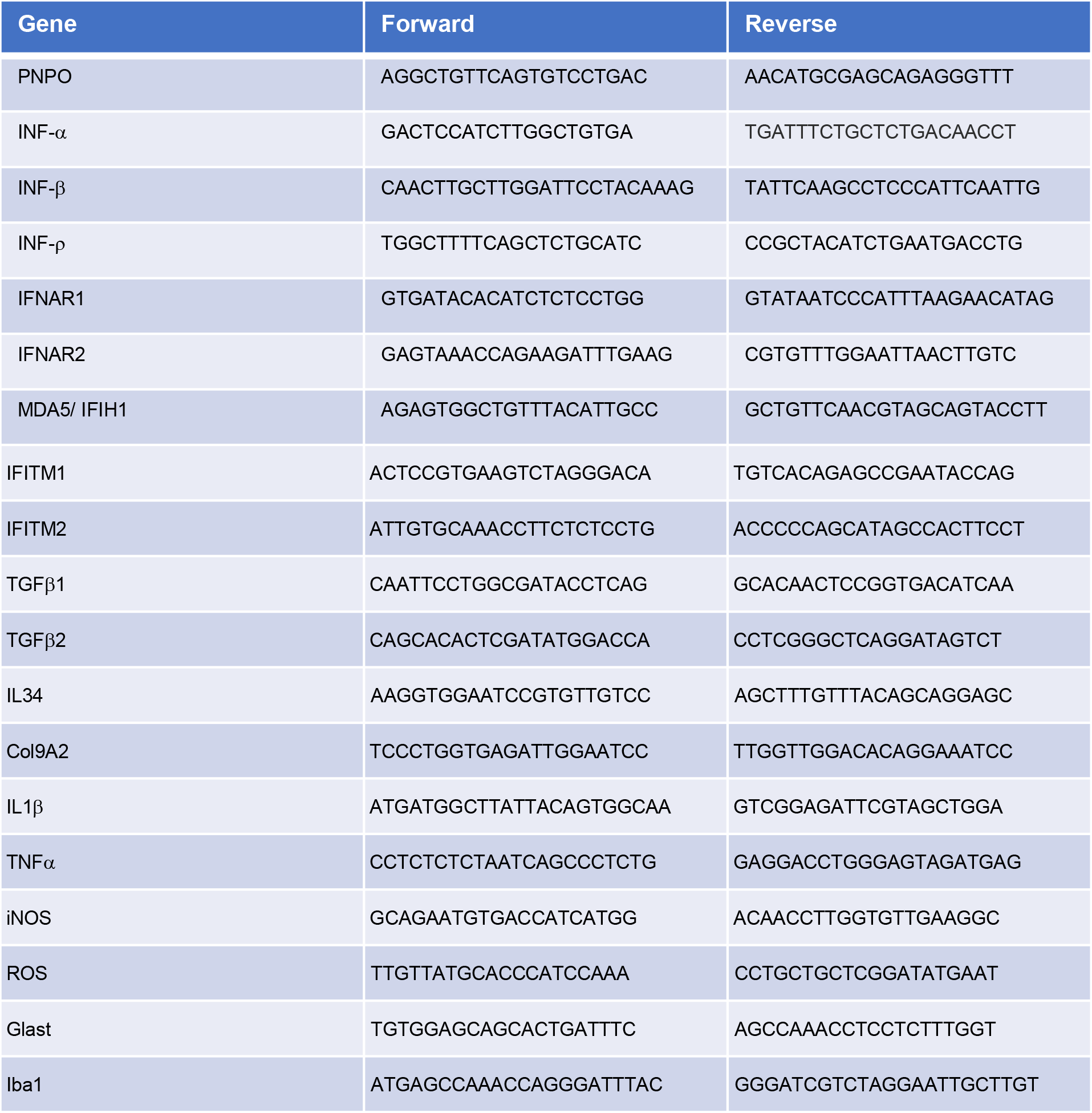
Primers used for RT-PCR study.

## Notes

### Competing Interest Statement

The authors have declared no competing interest.

